# Brainwide mapping of endogenous serotonergic transmission via chemogenetic-fMRI

**DOI:** 10.1101/122770

**Authors:** Andrea Giorgi, Sara Migliarini, Marta Gritti, Alberto Galbusera, Giacomo Maddaloni, Maria Antonietta De Luca, Raffaella Tonini, Alessandro Gozzi, Massimo Pasqualetti

## Abstract

Serotonergic transmission affects behaviours and neuro-physiological functions via the orchestrated recruitment of distributed neural systems. It is however unclear whether serotonin’s modulatory effect entails a global regulation of brainwide neural activity, or is relayed and encoded by a set of primary functional substrates. Here we combine DREADD-based chemogenetics and mouse fMRI, an approach we term “*chemo-fMRI”*, to causally probe the brainwide substrates modulated by phasic serotonergic activity. We describe the generation of a conditional knock-in mouse line that, crossed with serotonin-specific Cre-recombinase mice, allowed us to remotely stimulate serotonergic neurons during fMRI scans. We show that chemogenetic stimulation of the serotonin system does not affect global brain activity, but results in region-specific activation of a set of primary target regions encompassing parieto-cortical, hippocampal, and midbrain structures, as well as ventro-striatal components of the mesolimbic reward systems. Many of the activated regions also exhibit increased c-Fos immunostaining upon chemogenetic stimulation in freely-behaving mice, corroborating a neural origin for the observed functional signals. These results identify a set of regional substrates that act as primary functional targets of endogenous serotonergic stimulation, and establish causation between phasic activation of serotonergic neurons and regional fMRI signals. They further highlight a functional cross-talk between serotonin and mesolimbic dopamine systems hence providing a novel framework for understanding serotonin dependent functions and interpreting data obtained from human fMRI studies of serotonin modulating agents.

## Introduction

Serotonin (5-HT) is a modulatory transmitter produced by a small set of midbrain and brainstem nuclei which richly innervate forebrain target regions via long-range projections (Muzerelle *et al*, 2016). Such profuse innervation, along with the combined release of transmitters via pointwise synaptic signalling as well as non-synaptic ‘volume transmission’ (Descarries and Mechawar, 2000), can alter neuronal functions to define internal states that affect behavioural outputs in response to external stimuli. Recent research has shed light on the basic cellular and molecular machinery engaged by 5-HT (Lesch and Waider, 2012). However, many question regarding the systems-level substrates modulated by 5-HT transmission remain unanswered. For one, it is unclear whether serotonin’s modulatory action entail a global regulation of brainwide neural activity, or it is relayed and encoded by a set of primary functional targets.

Neuroimaging studies have employed pharmacological manipulations to probe the brainwide targets of 5-HT. The vast majority of these studies have primarily focused on the use of functional magnetic resonance imaging (fMRI) as a proxy for the neuronal activity directly elicited by pharmacological agents, or by mapping second-order modulatory effects on task-elicited fMRI responses (Anderson *et al*, 2008). This approach, termed pharmacological fMRI, has also been back-translated to animal studies (Gozzi *et al*, 2012; Klomp *et al*, 2012). However, while useful in identifying possible brainwide substrates of neuromodulatory action, drug-based approaches may be contaminated by off-target receptor contributions (Ofek *et al*, 2012), and typically engage subtype-specific receptors systems that may not be representative of the overall neuromodulatory action of a transmitter systems. Furthermore, drug-elicited signals may contain peripheral vasotonic or cerebrovascular contributions that cannot be easily disentangled from the drug’s primary neural effects (Martin and Sibson, 2008). This aspect is especially relevant when hemodynamic measures of brain function are used, given that 5-HT can perivascularly modulate blood flow, independent of its central effects (Cohen *et al*, 1996). As a result, it remains unclear where and how phasic serotonergic activity specifically influences regional or global brain functional activity, and the corresponding fMRI responses.

The advent of light-based or chemically-mediated cell-manipulation methods has made possible to causally link the activity of focal neuronal populations with specific circuital and behavioural outputs (Park and Carmel, 2016). When combined with *non invasive* neuroimaging methods, cell-type specific manipulations can be used to map the functional substrates of endogenous modulatory transmission, without the confounding contribution of peripherally-elicited vasoactive or pharmacological effects (Ferenczi *et al*, 2016). Towards this goal, here we describe the combined use of mouse cerebral-blood volume based fMRI (Galbusera *et al*, 2017) and cell-type specific DREADD (Designed Receptors Exclusively Activated by Designed Drugs (DREADDs) (Armbruster *et al*, 2007)) chemogenetics, an approach we term ‘chemo-fMRI’, to map the brainwide functional targets of phasic 5-HT stimulation.

To allow for stable and reproducible phasic 5-HT stimulation across animals we generated a conditional knock-in mouse that we crossed with Pet1 Cre-transgenic mice (Pelosi *et al*, 2014), permitting to remotely activate 5-HT-producing neurons during fMRI scans with reduced inter-subject variability and without the need of invasive surgery. Our results show that chemogenetic 5-HT stimulation does not affect global brain activity, but results in region specific activation of a set of primary target regions encompassing parieto-cortical, hippocampal, and midbrain structures, as well as ventro-striatal components of the mesolimbic reward systems. Importantly, we also show that pharmacological boosting of 5-HT levels via administration of the selective serotonin reuptake inhibitor citalopram produces widespread brain deactivation, a finding plausibly reflecting a contribution of peripheral vasoconstrictive effect of 5-HT on blood vessels (Cohen *et al*, 1996) and underscoring the need to carefully control for possible vascular contributions in functional brain mapping of serotonergic drugs. Collectively, our results reveal a set of regional substrates that act as primary functional targets of endogenous serotonergic stimulation and establish causation between phasic activation of 5-HT neurons and regional fMRI signals. They also provide a novel framework for understanding 5-HT-dependent functions and interpreting data obtained from human fMRI studies of 5-HT modulating agents.

## Results

### hM3Dq-mediated modulation of serotonergic neurons

To enable stable pan-neuronal stimulation of 5-HT producing cells, we first generated a hM3Dq conditional knock-in mouse encompassing the integration of hM3Dq-mCherry double-floxed inverse open reading frame (DIO-hM3Dq) within the ROSA26 genomic locus (Fig. 1). In the designed mouse line, stable Cre-mediated somatic recombination is required for conditional transcriptional activation (CAG promoter-driven) of the hM3Dq gene, whose expression could be probed by the presence of in-frame-fused mCherry reporter (Fig. 1). To enable hM3Dq expression in 5-HT producing neurons, DIO-hM3Dq mice were crossed with the Pet1_210_-Cre transgenic mouse line (Pelosi *et al*, 2014) to generate hM3Dq/Pet1-Cre mice. To corroborate and quantify the frequency and specificity of hM3Dq expression in 5-HT producing neurons, we performed double immunofluorescence analysis on triple trans-heterozygous hM3Dq/Pet1-Cre/Tph2^GFP+/-^ mice. Tph2^GFP+/-^ express GFP under the control of the endogenous promoter of the 5-HT marker gene Tph2, such to enhance the sensitivity of 5-HT-based cell labelling (Migliarini *et al*, 2013). These analyses revealed that hM3Dq is expressed in the vast majority (95%) of 5-HT producing neurons (n=3 mice, 2909±119.64 counted cells, 2658.33±119 mCherry(+) eGFP(+) cells, 124.33±12.42 mCherry (-) eGFP(+) cells, Fig. 2A).

**Figure 1.**
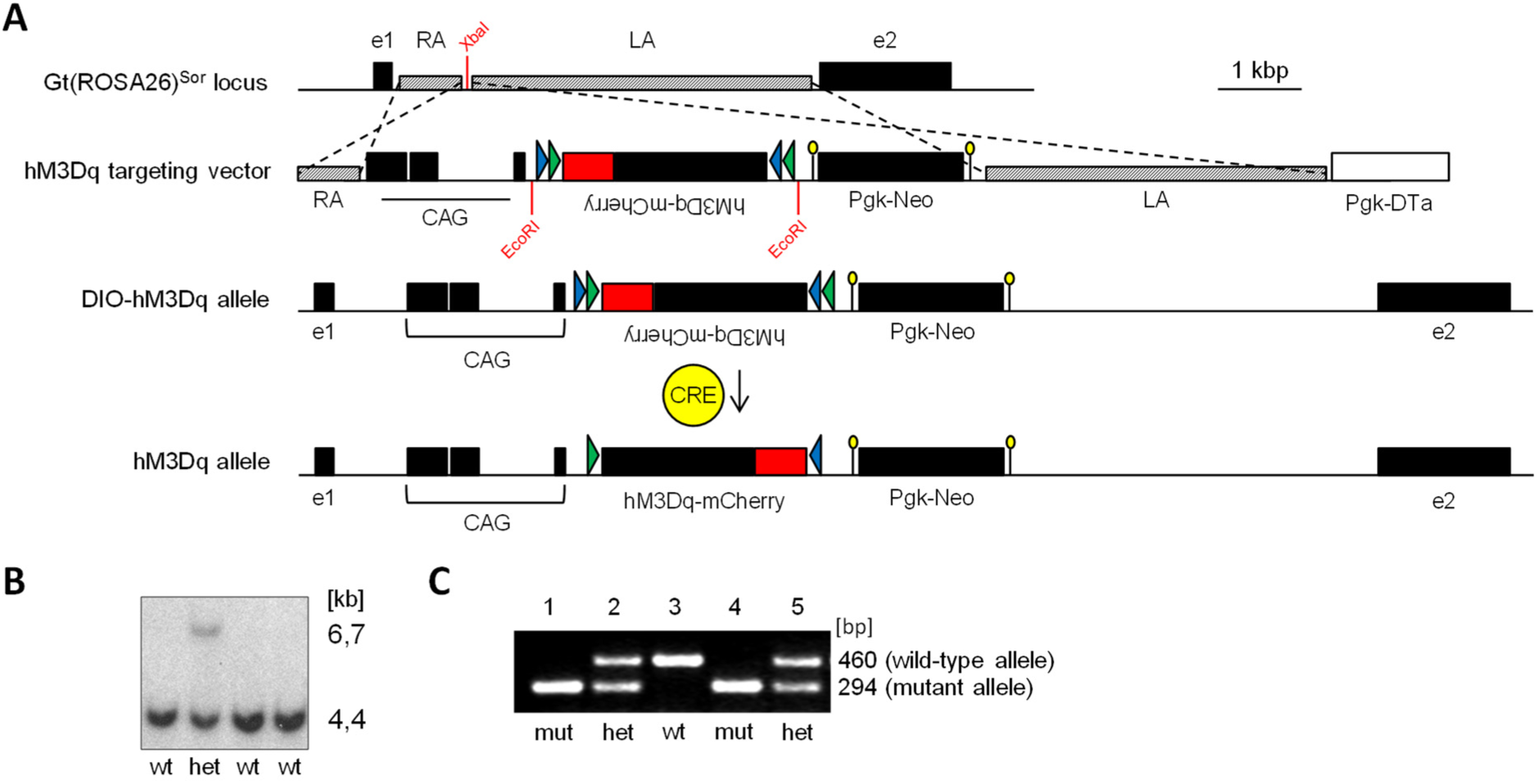
Generation of hM3Dq conditional knock-in mouse line. Targeted integration of the hM3Dq construct in the ROSA26 genomic locus. (A) From top to the bottom: wild-type organization of ROSA26 mouse genomic locus; hM3Dq targeting vector, the red box symbolizes the in-frame-fused mCherry reporter, green (LoxP) and blue (Lox2722) arrowheads indicate the location of Lox sites, yellow circles are used for FRT sites; transcriptionally inactive DIO-hM3Dq allele, in which the hM3Dq-mCherry coding sequence (CDS) is inverted with respect to the CAG promoter; the transcriptionally active hM3Dq allele, in which the hM3Dq-mCherry CDS was reverted by a Cre-mediated somatic recombination. (B) Southern blot analysis on genomic DNA extracted from four embryonic stem cells clones after *HindIII* digestion (hM3Dq allele=6,7 kb; wild-type allele=4,4 kb). (C) PCR amplification of ROSA26 locus from DNA extracted from a wild-type (3), two mutants (1 and 4) and two heterozygous (2 and 5) DIO-hM3Dq mice.

**Figure 2.**
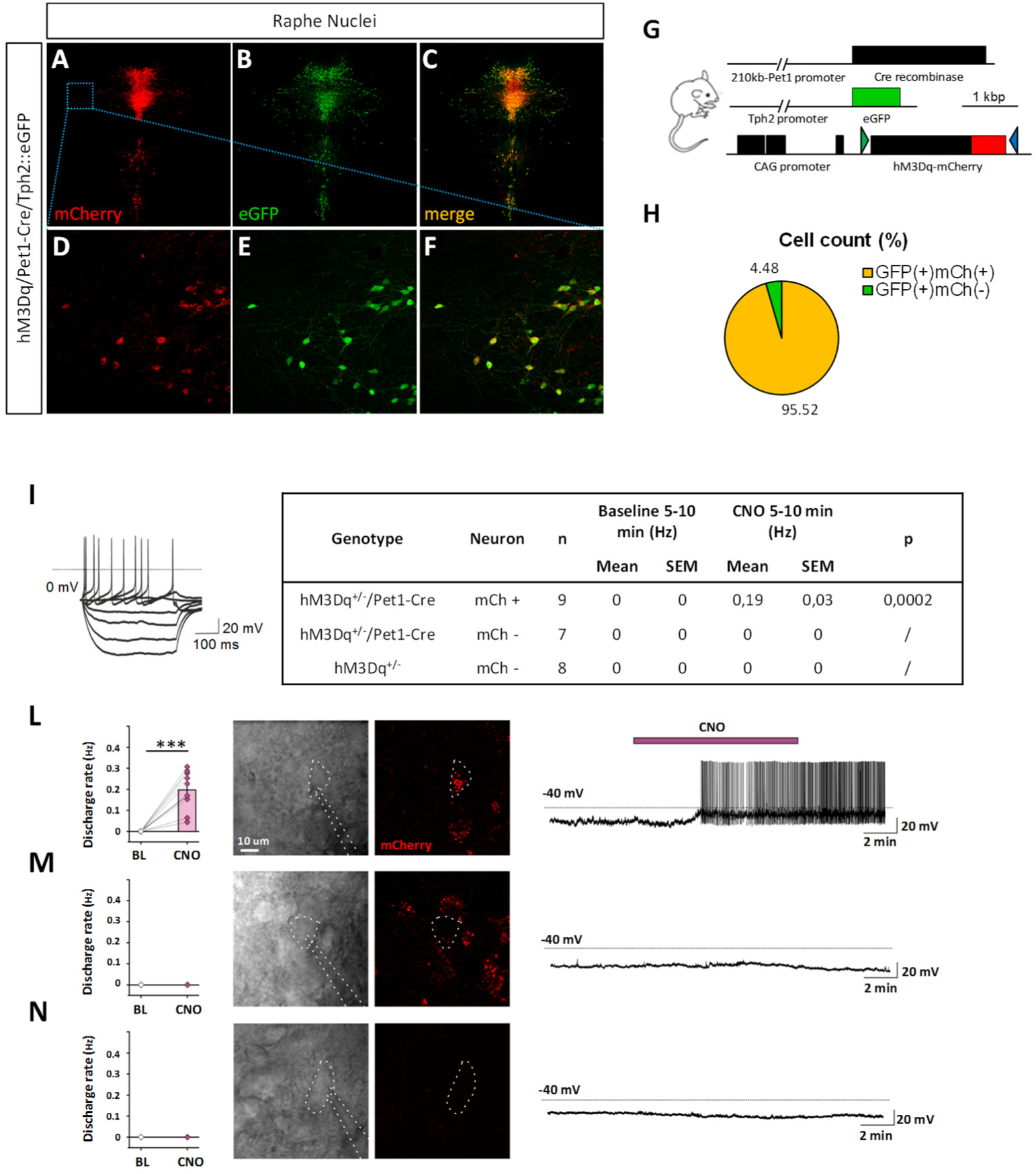
Selective hM3Dq expression in 5-HT neurons and electrophysiological validation. (A-F) Double immunohistochemical assay showing (A,D) mCherry expression, (B,E) GFP expression (Tph2) and (C,F) merged channels in the DRN (A-C) and DRN lateral wings (D-F) in hM3Dq/Pet1-Cre/Tph2^GFP+/-^ mice. (G) Schematic genotype representation of hM3Dq/Pet1-Cre/Tph2^GFP^+^/-^ triple heterozygous mice used for immunohistochemical labelling, green (LoxP) and blue (Lox2722) arrowheads indicate the location of Lox sites after Cre-mediated recombination. (H) Quantification of hM3Dq-expressing 5-HT neurons: mCherry (in-frame-fused with hM3Dq) was found to be present in the 95% of the total eGFP-positive (Tph2-positive) cells (I) Electrophysiological properties defining 5-HT neurons (L) Discharge rate, image of the recorded cell and single-cell firing tracking of CNO treated hM3Dq/Pet1-Cre mCherry(+) neuron. (M) Discharge rate, image of the recorded cell and single-cell firing tracking of CNO treated hM3Dq/Pet1-Cre mCherry(-) neuron. (N) Discharge rate, image of the recorded cell and single-cell firing tracking of CNO treated wild-type 5-HT neuron. The purple line indicates CNO administration time window.

We next recorded the electrophysiological response induced by CNO administration in hM3Dq-expressing 5-HT neurons via whole-cell patch clamp recordings in coronal brain slices encompassing the dorsal raphe nucleus (DRN), the main anatomical source of forebrain-innervating 5-HT neurons. We confirmed serotonergic identity of patched cells based on their intrinsic electrophysiological properties including lack of time-dependent depolarization in response to hyperpolarizing current pulses (Calizo *et al*, 2011). As expected, mCherry-positive neurons in hM3Dq/Pet1-Cre mice were robustly activated by CNO, resulting in sustained firing rates (Fig.2, B). We did not observe any CNO-induced firing rate alterations in any of the neighboring mCherry-negative neurons probed (n=8), or in 5-HT neurons in DIO-hM3Dq control mice (n=7) (Fig.3-tracks). These results suggest that hM3Dq/Pet1Cre mice can reliably be employed to endogenously stimulate 5-HT neuronal firing upon CNO administration.

**Figure 3.**
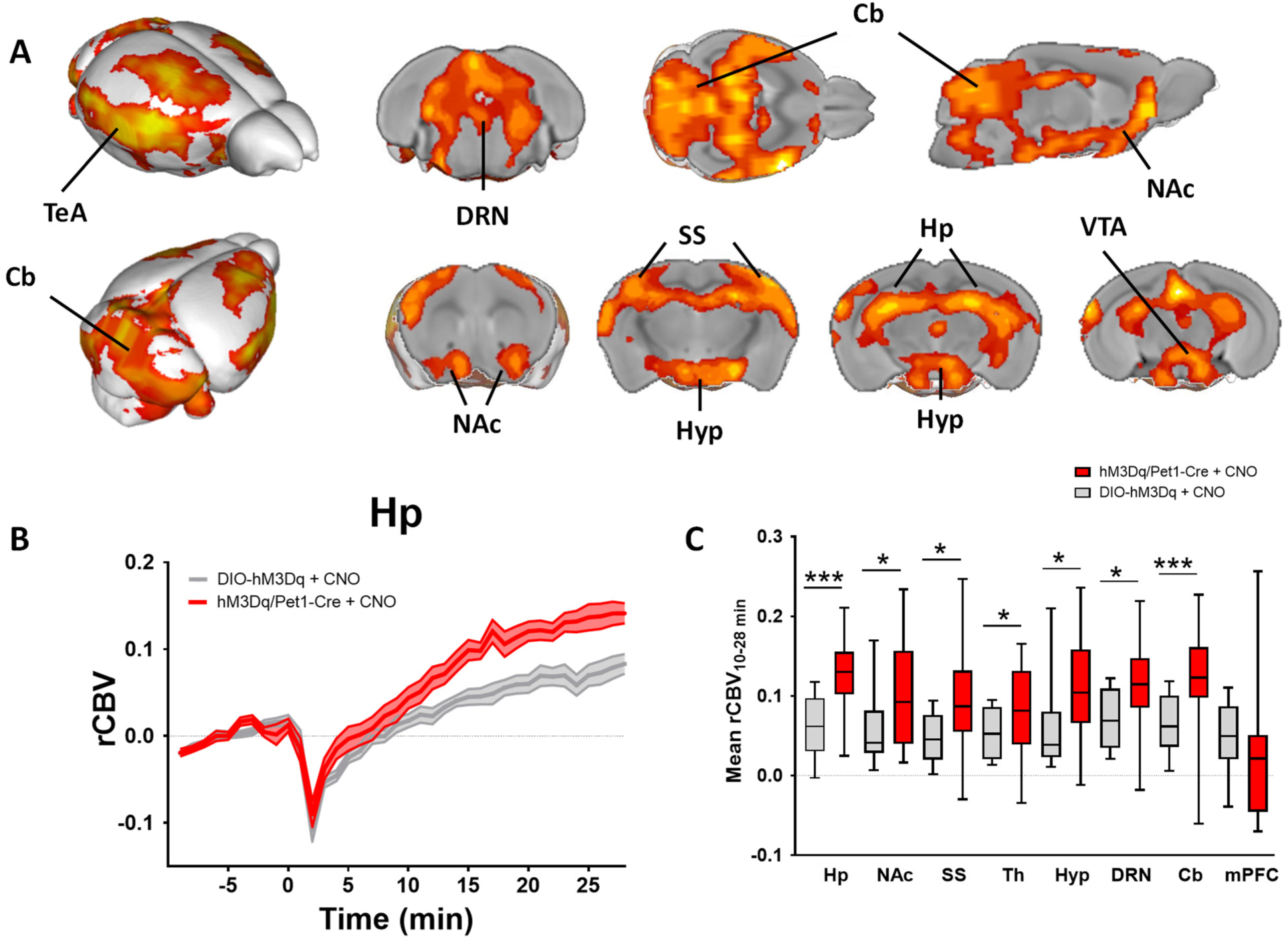
fMRI response to chemogenetic stimulation of 5-HT-producing neurons. fMRI activation produced by intravenous CNO administration (0.5 mg/kg, iv) in the brain of hM3Dq/Pet1-Cre mice (n=20) vs CNO-treated DIO-hM3Dq controls (n=14). (A) Three-dimensional rendering of the anatomical distribution of the mouse brain regions significantly activated by acute CNO administration vs controls (Z>2, cluster corrected at p=0.05) (B) Illustrative fMRI timecourse in the hippocampus, CNO was administered at time 0. (C) Boxplot representation of region-specific rCBV changes. Data are plotted within each group as individual means calculated from the minute 10 to the minute 28 post CNO administration (*p<0.05, **p< 0.01, Student t test, false discovery rate correction q=0.05). Hp: hippocampus; NAc: nucleus accumbens; SS: somatosensory cortex; Th: thalamus; Hyp: hypothalamus; DRN: dorsal raphe nucleus; Cb: cerebellum, mPFC: prefrontal cortex.

### Chemo-fMRI mapping of endogenous 5-HT neurotransmission

To determine the feasibility of using fMRI to map cell-type specific DREADD-based neural responses, we first performed a dose-response titration (0.5-1-2 mg/kg i.v.) of the fMRI response produced by CNO *per se* in wild-type animals devoid of hM3Dq receptors. Intravenous CNO administration (1 and 2 mg/kg i.v.) produced robust relative cerebral blood volume (rCBV) increases in most of the examined volumes of interest (Fig. S2). At a dose of 0.5 mg/kg i.v., CNO-induced rCBV alterations were qualitatively distinguishable from reference baseline signal (vehicle) only in hypothalamic and hippocampal areas (Fig. S2). These effects were recapitulated by regional quantification of mean rCBV in volumes of interest (Fig. S3), which highlighted robust increases in rCBV baseline in most of the examined anatomical regions at dose of 1 and 2 mg/kg, and negligible effects at a 0.5 mg/kg dose (Fig. S3, p<0.05, Student t test, FDR corrected, q=0.05). These results suggest that high doses of CNO can elicit unspecific fMRI signal changes. To better control for spurious CNO-induced fMRI signal, we performed chemo-fMRI mapping by administering a low-dose of CNO (0.5 mg/kg) to both hM3Dq/Pet1-Cre *and* control DIO-hM3Dq mice, and using the latter cohort as baseline reference control for quantification and statistical mapping of elicited responses.

Chemogenetic activation of 5-HT producing neurons in hM3Dq/Pet1-Cre mice elicited robust region-specific rCBV increases in several cortical and non-cortical substrates with respect to DIO-hM3Dq CNO-treated controls (Fig. 3). The observed fMRI timecourses were characterized by a slow but gradual onset with a clear differentiation with baseline reference signal in DIO-hM3Dq subject occurring between 10-15 min after CNO injection (Fig. S5). Voxelwise mapping of the elicited chemo-fMRI response revealed a pattern of increased rCBV encompassing parietal cortical regions such as somatosensory and motor cortices, as well posterior insular and temporal association regions (Fig. 3, Z>2, cluster corrected at p=0.05). We also observed robust rCBV increases in several subcortical substrates, including hippocampal areas, the hypothalamus, the dorsal raphe and cerebellum. A prominent involvement of ventral tegmental area and its mesolimbic terminals in the ventral striatum and nucleus accumbens was also apparent. Importantly, CNO administration did not result in major or abrupt blood pressure alterations at any of the doses tested in any of the experimental groups examined (Fig. S4).

To corroborate a neural origin of hM3Dq-mediated rCBV increases, we performed c-Fos immunofluorescence mapping in a separate cohort of freely behaving in hM3Dq/Pet1-Cre and control DIO-hM3Dq mice. We observed robust region-specific increase of c-Fos expressing neurons in many of the substrates identified with fMRI, including nucleus accumbens, hippocampus, anterior thalamus, hypothalamic nuclei and raphe nuclei of hM3Dq/Pet1-Cre mice with respect to control animals (Fig. 4, p<0.05, all regions, Student t test, FDR q=0.05). No c-Fos induction was instead observed in the cerebellum and somatosensory cortex, two regions exhibiting significant rCBV responses in chemo-fMRI mapping.

**Figure 4.**
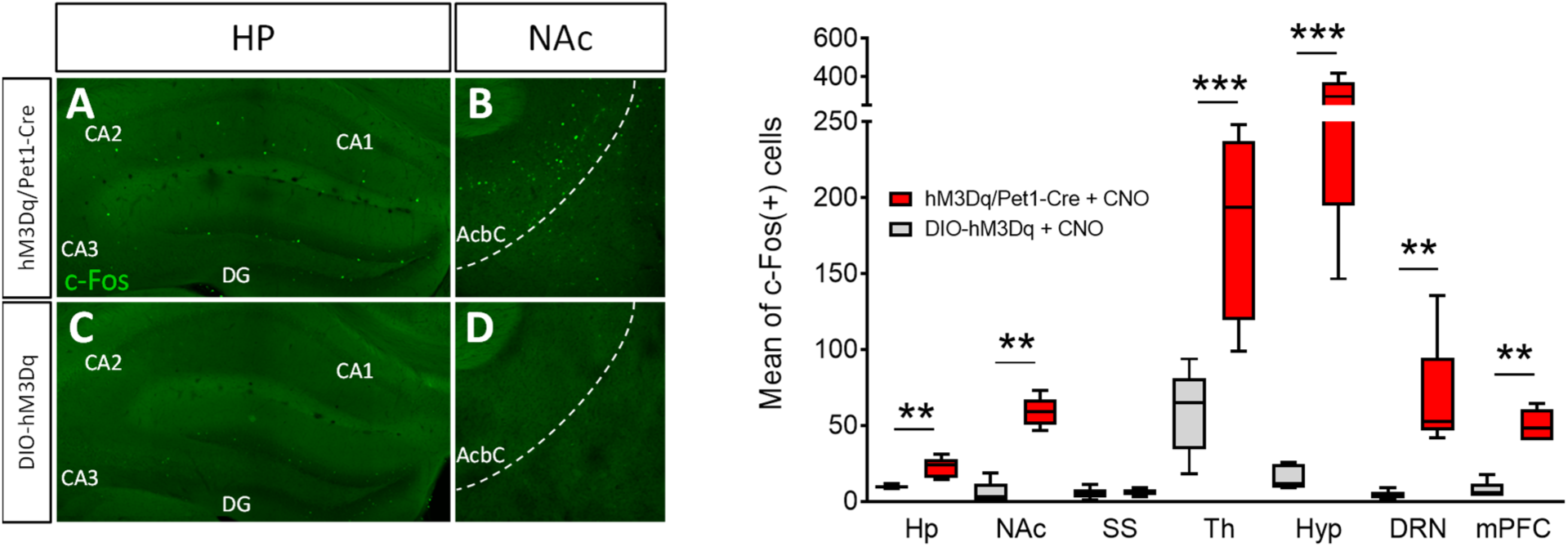
c-Fos induction produced by hM3Dq-mediated phasic activation of 5-HT neurons. Regional c-Fos immunofluorescence in the brain of hM3Dq/Pet1-Cre mice (n=5) and control (DIO-hM3Dq) subjects treated with CNO (n=5). (A-D) Distribution of c-Fos-positive (green) cells (A) in the hippocampus and (B) nucleus accumbens of hM3Dq/Pet1-Cre and control mice (C, D). (E) Quantification of c-Fos-positive cells in the regions of interest (*p<0.05, **p< 0.01, Student t test, false discovery rate correction q=0.05). Hp: hippocampus; NAc: nucleus accumbens; SS: somatosensory cortex; Th: thalamus; Hyp: hypothalamus; DRN: dorsal raphe nucleus; Cb: cerebellum; mPFC: prefrontal cortex; CA1-2-3: cornu ammonis 1-2-3; DG: dentate gyrus AcbS: nucleus accumbens shell.

### Systemic 5-HT reuptake inhibition produces fMRI deactivation

An underlying assumption of our work is that the signals elicited by CNO in hM3Dq/Pet1-Cre mice primarily reflect central 5-HT neuronal stimulation, which in turns elicits fMRI signals via neurovascular coupling. By contrast, fMRI mapping of systemically-administered 5-HT ligands may be contaminated or masked by direct stimulation of endothelial 5-HT receptors, leading to peripherally-mediated vascular responses which in the case of 5-HT are typically vasoconstrictive (Cohen *et al*,1996). To investigate this hypothesis, we carried out fMRI mapping upon systemic administration of the selective 5-HT reuptake inhibitor citalopram (5-10 mg/kg), a drug leading to increased peripheral and central 5-Ht levels (Blardi *et al*, 2005; Calcagno *et al*, 2007). Interestingly, citalopram (10 mg/kg. iv.) produced widespread foci of rCBV signal decrease in a widespread set of cortical and subcortical regions, as seen with voxelwise mapping (Fig. 5, Z >1.8, cluster corrected p=0.05). Although the effect was widely distributed, the deactivation was particularly prominent in the prefrontal cortex (Fig. 5B, Fig. S6). Citalopram administration was not associated with appreciable blood pressure alteration at any of the doses tested (Fig. S4).

**Figure 5.**
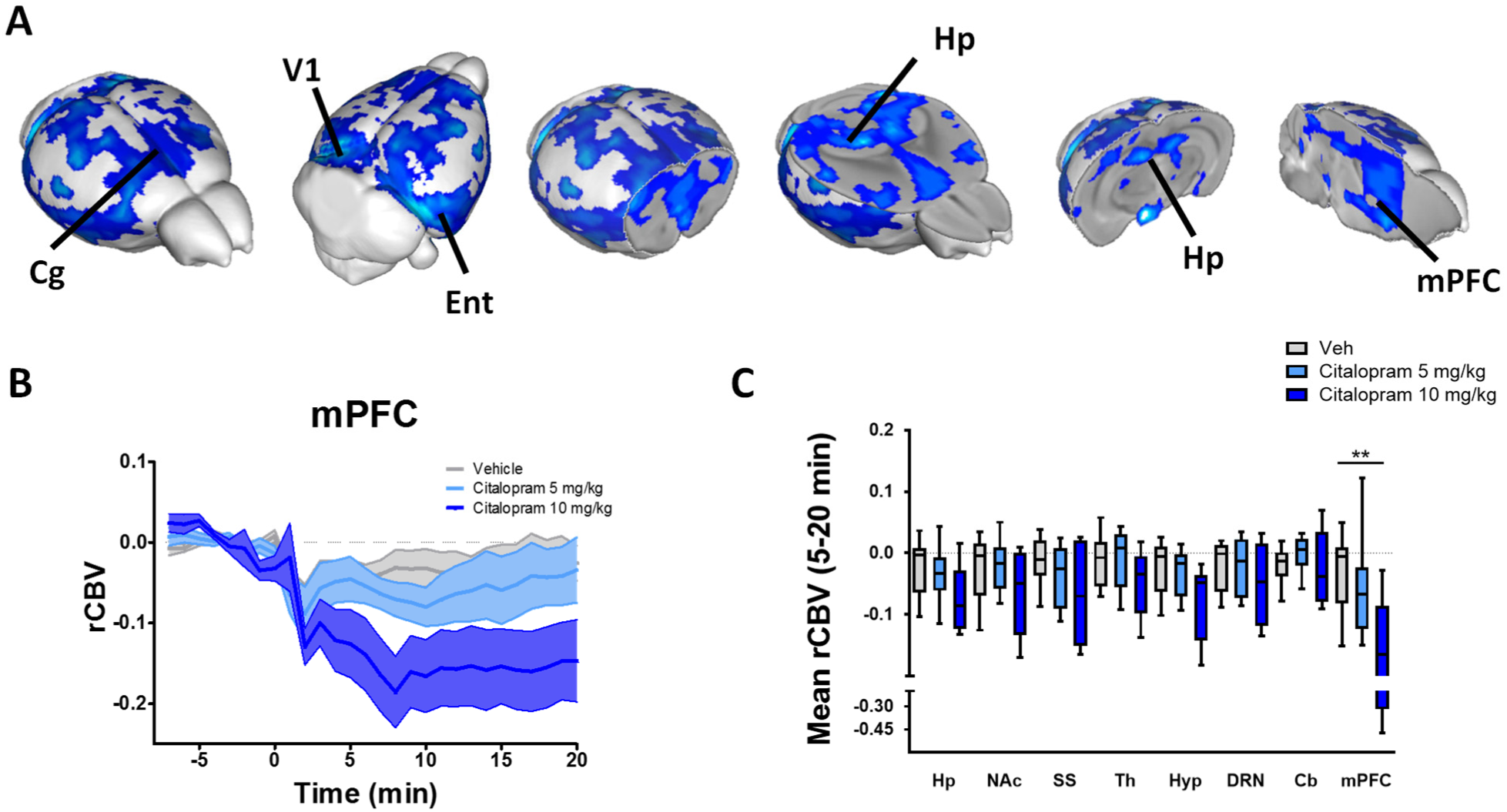
Brainwide fMRI response to systemic citalopram administration. (A) 3D rendering of the anatomical distribution of the mouse brain regions significantly affected by acute citalopram (10 mg/kg) (n=7) vs vehicle (z>1.8, cluster correction p=0.05) (n=10). (B) Representative temporal profile of CNO-induced rCBV response in the prefrontal cortex of mice treated with citalopram 10 mg/kg vs controls. Citalopram or vehicle were administered at time 0. (C) rCBV quantification in representative volumes of interest (*p<0.05, **p< 0.01, Student t test, false discovery rate q=0.05). Hp: hippocampus; NAc: nucleus accumbens; SS: somatosensory cortex; Th: thalamus; Hyp: hypothalamus; DRN: dorsal raphe nucleus; Cb: cerebellum, mPFC: prefrontal cortex; Cg: cingulate cortex; V1: Visual cortex; Ent: entorhinal cortex.

## Discussion

Serotonin is an archetypical neruomodulatory monoamine characterized by restricted neural localization but with broad and far-ranging modulatory properties. Despite extensive research, the macroscale functional substrates endogenously modulated by 5-HT in the intact brain remain elusive. By using chemo-fMRI, we describe the primary brainwide targets of phasic serotonergic stimulation, and establish causation between phasic activation of 5-HT neurons and regional fMRI signals. Our results provide a novel framework for understanding 5-HT-dependent function and interpreting data obtained from human fMRI studies of 5-HT modulatory agents.

While the density of 5-HT ascending projection is highly variable across neuroanatomical regions, virtually no area in the central nervous system is devoid of 5-HT innervation (Muzerelle *et al*, 2016). The observation of region-specific responses in spite of the use of drug-like pan-serotonergic stimulation provides evidence that phasic serotonergic neurotransmission engages a set of neuroanatomical targets serving as primary, fast-responding effectors of phasic release of this neurotransmitter. This notion is consistent with a role of phasic 5-HT transmission as a mediator of strongly-arousing species-normative consummatory behaviours that strongly engage evolutionary-ancient ventral limbic substrates, and that could involve a the excitatory contribution of glutamate co-release (El Mestikawy *et al*, 2011; Ferrari *et al*, 2003; Liu *et al*, 2014; Takahashi *et al*, 2012). Among the mapped regions, the presence of parietal cortices is in keeping with early metabolic and cerebral blood flow-based mapping upon electrical or chemical stimulation of raphe regions (Cohen *et al*, 1996; Cudennec *et al*, 1987; Underwood *et al*, 1995), and the robust activation of hippocampal regions is consistent with the especially dense 5-HT innervation of these areas (Migliarini *et al*, 2013). Importantly, our mapping provides functional evidence that serotonergic activation can also modulate the activity of mesolimbic dopamine regions, such as the nucleus accumbens. This finding strengthens evidence of a tight functional interplay between these two key modulatory systems, and is in good agreement with the emerging view of role of 5-HT as a modulator of reward processing and mesolimbic dopamine substrates (Li *et al*, 2016; Luo *et al*, 2015). Interestingly, recent mapping of chemogenetically-stimulated neurons via 2-fluoro-dexoy-glucose PET did not highlight cortico-hippocampal modulation of brain activity, but revealed small foci of reduced metabolism in thalamic areas (Urban *et al*, 2016). Several experimental factors could account for these discrepant results, including different spatiotemporal sensitivity of fMRI with respect to PET these measurements, plus the contribution of unspecific effects of CNO in the study from Urban and colleagues, which did not employ a control CNO-treated group of subjects.

Our findings also provide a novel reference framework for the interpretation of fMRI imaging studies employing 5-HT-targeting agents. In this respect, the divergent functional effects observed with chemogenetic stimulation and the 5-HT reuptake inhibitor citalopram is especially relevant, as it suggests that central stimulation and systemic 5-HT level boosting can lead to opposing hemodynamic responses, a contribution that needs to be taken into account when hemodynamic readouts are used as surrogates for 5-HT induced neural manipulations. A role for 5-HT in controlling central and perivascular control of vascular activity has been long established, with evidence of a dominating vasoconstrictive action of this neurotransmitter via 5-HT_1A_ and possibly 5-HT_2A_ receptors located in brain and peripheral vascular cells (reviewed by Cohen *et al*, 1996). In keeping with this, pharmacological stimulation of 5-HT_1A_ receptors has been reliably shown to induce widespread microvascular rCBV decreases (Gozzi *et al*, 2010; Mueggler *et al*, 2011). Because 5-HT reuptake inhibition rapidly increases circulating levels of 5-HT (Blardi *et al*, 2005; Calcagno *et al*, 2007; Zolkowska *et al*, 2008), and systemic 5-HT administration produces peripherally-driven arterial vasoconstriction (Dieguez *et al*, 1981; Toda and Fujita, 1973), citalopram-induced rCBV decreases are likely to reflect a dominant peripheral vasoactive contribution of circulating 5-HT. Interestingly, prior blood oxygen level-dependent (BOLD) fMRI mapping of citalopram in rats revealed increased cortical fMRI responses that appear to be at odds with this hypothesis (Sekar *et al*, 2011). A number of methodological discrepancies could account for these inconsistent results, the use of deeper anesthesia levels by Sekar et al., as well as the use of BOLD fMRI, as opposed to rCBV like in the present work, being the two most notable ones. With respect to the employed fMRI readout, it should be emphasized that unlike BOLD, rCBV is a direct and reliable indicator of microvascular activity, and appears to be characterised by a more direct relationship with underlying neuronal activity with respect to traditional BOLD fMRI (Schridde *et al*, 2008). Further chemo-fMRI studies of 5-HT function employing BOLD fMRI are warranted to conciliate these experimental discrepancies.

The observation of significant c-Fos induction in many of the regions exhibiting significant 5-HT induced chemo-fMRI activation is consistent with a neural origin of the mapped functional signals, and the presence of minimal interference of the light anesthesia used in functional mapping. Given the independent nature of cellular (i.e., c-Fos induction) and hemodynamic (i.e., phMRI) measures of brain function, these correspondences argue against spurious contributions arising from a putative direct vasoactive action of CNO. Discrepancies between imaging and molecular data are also to be noted especially in cortical regions and in the cerebellum. Such discrepancies are not surprising given the diverse neuro-physiological mechanisms underlying these two experimental readouts. Indeed, hemodynamic responses are generally thought to reflect local synaptic input (Logothetis *et al*, 2001), whereas the c-Fos protein induction is an indirect marker of cellular activation that is not elicited in all CNS cell groups (Cirelli and Tononi, 2000). In keeping with this, previous studies showed that c-Fos induction does not always correspond to activation shown by metabolic or hemodynamic-based brain mapping (Gass *et al*, 1997; Gozzi *et al*, 2012; Stark *et al*, 2006).

Interestingly, our functional mapping did not highlight phasic functional involvement of brain substrates known to be innervated by 5-HT neurons, such as the medial prefrontal cortex (Puig and Gulledge, 2011) or the caudate putamen (Fadda *et al*, 2005). This observation is unlikely to reflect the lack of hM3Dq-mediated stimulation in these regions, given the almost complete transduction of DREADD receptors in 5-HT containing raphe nuclei, as well as evidence that striatal 5-HT transmission can be successfully manipulated using an analogous knock-in intersectional strategy using Pet1 Cre mice (Raffaella Tonini, personal communication). The use of light anaesthesia is also unlikely to result in regional suppression of 5-HT responses in these regions, as the regimen employed here preserves the functional organization of the mouse brain (Gozzi and Schwarz, 2015), and has been successfully employed to map the response of monoaminergic agents (Gozzi *et al*, 2012; Gozzi *et al*, 2013). A differential contribution of wiring and volume transmission (Descarries *et al*, 2000), as well as regional combinations of receptor subtypes characterized by different transductional signalling or cell-type specific distribution serve as possible explanation for the regional effects mapped, and the phasic functional engagement of a selective subset of brain targets upon endogenous 5-HT stimulation.

From a methodological standpoint, this work represents to our knowledge the first demonstration of the combined use of chemogenetic and fMRI to unravel the large-scale substrates modulated by focal neuronal stimulations. Our approach follows analogous attempts to combine chemogenetics with non-invasive mouse brain imaging, such as metabolic mapping via positron emission tomography (DREAMM - Urban *et al*, 2016). Our data demonstrate that the ensuing chemo-fMRI responses exhibit neuronal specificity, are sustained, and can be employed to obtain to non-invasive map regional manipulations in the living mouse brain. This approach nicely complements current opto-fMRI simulations (Lee *et al*, 2010) by permitting brainwide functional mapping without the need of invasive cranial probes, and the confounding contribution of heat-induced vasodilation (Rungta *et al*, 2017). As such, chemo-fMRI appears to be optimally suited to the investigation of drug-like sustained modulatory stimulation/inhibition, the deconstruction or manipulation of steady-state network activity via fMRI connectivity mapping (Gozzi *et al*, 2015; Sforazzini *et al*, 2014). While here employed to study pan-neuronal 5-HT modulation, this approach can be straightforwardly expanded to manipulate focal circuits and neuronal ensembles via intersectional genetics or retrograde viral vectors.

In conclusion, we describe the use of chemo-fMRI to map the brainwide substrates of modulatory transmission in the intact mammalian brain. We show that chemogenetic stimulation of endogenous serotonin neurons results in regionally-specific activation of a set of cortical and subcortical targets that serve as primary functional mediators of the phasic effect of 5-HT transmission. We also show that phasic activation of serotonergic neurons and systemic 5-HT reuptake inhibition can produce opposing hemodynamic effects. Our findings provide a novel framework for understanding serotonin dependent functions, and interpreting data obtained from human fMRI studies of serotonin modulating agents.

## Acknowledgements

This work was supported by Italian Ministry of Education, University and Research (MIUR) (Prin 2008, 200894SYW2), Toscana Life Sciences Foundation (Orphan_0108 program) and Norwegian Research Council to M. P. G.M. was supported by PhD program from University of Pisa. S.M. was supported by Regional Program and European Social Fund. A.G. received funding from the Simons Foundation (SFARI 314688 and 400101) and Brain and Behaviour Foundation (NARSAD). We thank Bryan Roth for generously supplying hM3Dq and hM3Di cDNA, Carola Canella for helping with figure generation, C. Valente for his excellent technical assistance and members of our laboratories for valuable discussions and comments on the manuscript.

---

**Figure S1.**
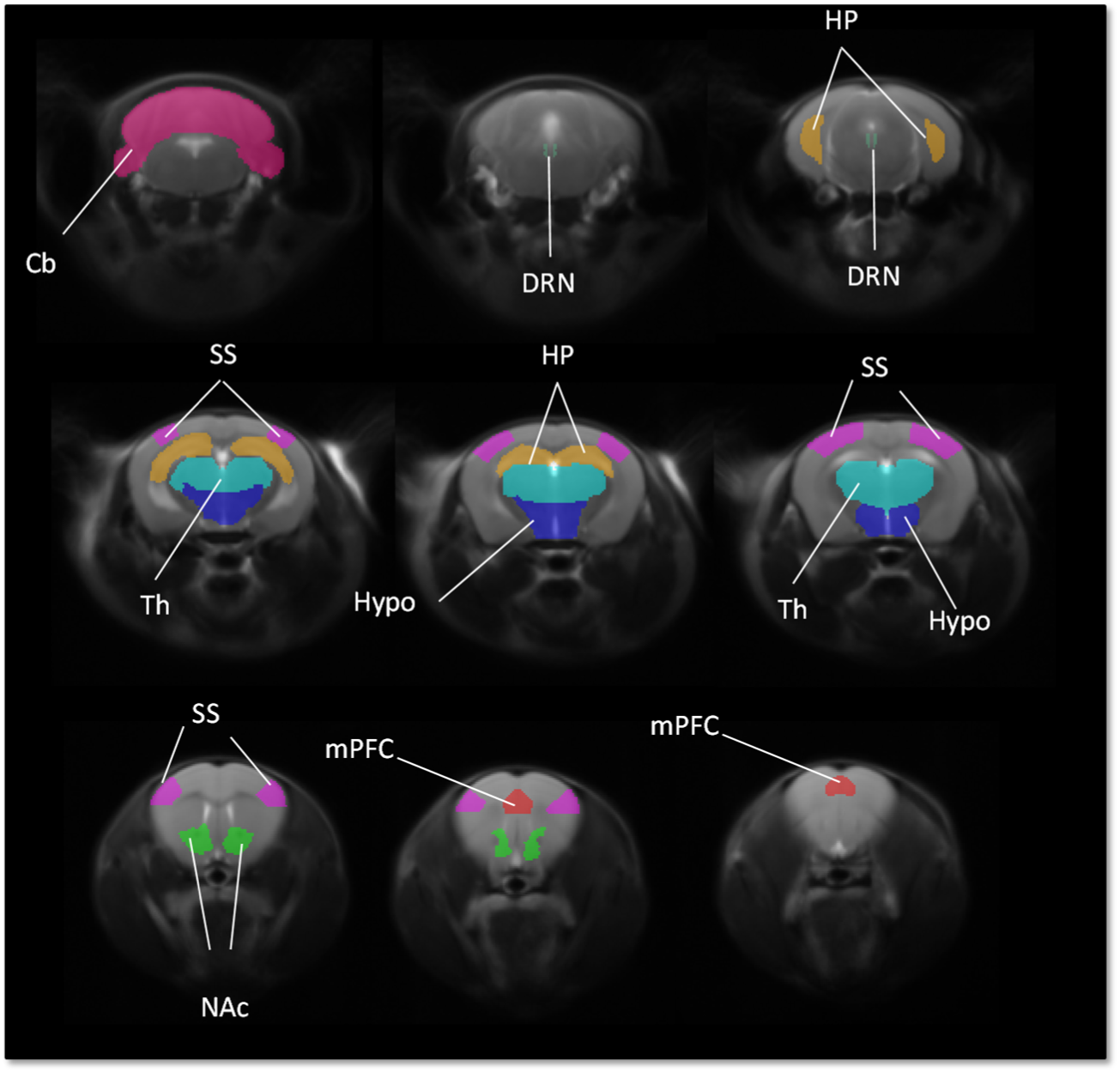
Volumes of interest used for rCBV quantification. Hp: hippocampus; mPFC: prefrontal cortex; DRN: dorsal raphe nucleus; NAc: nucleus accumbens; Th: thalamus; Hyp: hypothalamus; SS: somatosensory cortex; Cb: cerebellum.

**Figure S2.**
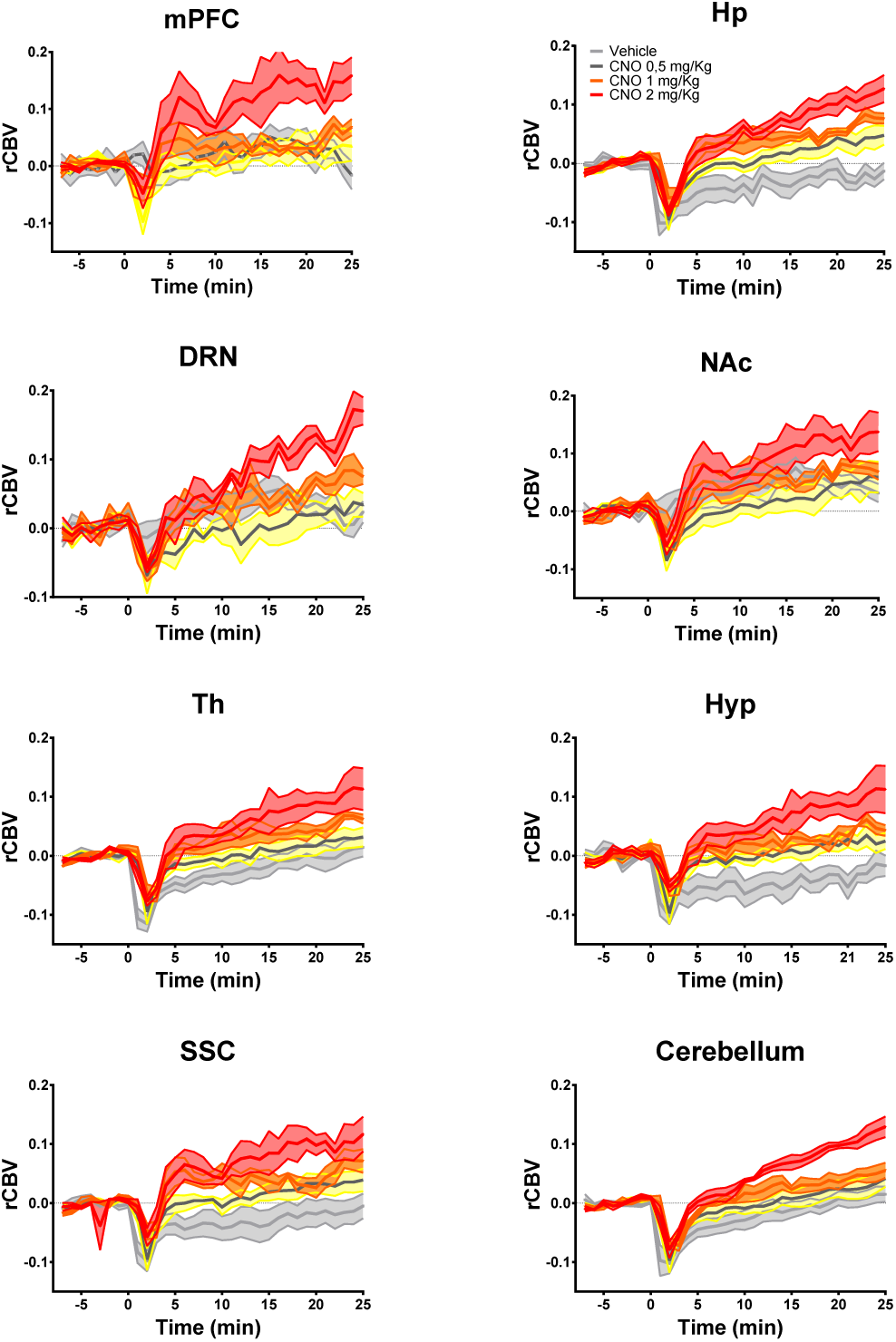
rCBV response produced by intravenous CNO administration in wild-type mice. CNO was administered intravenously at doses of 0.5 (n=6), 1 (n=4) or 2 mg/kg (n=5). rCBV timecourses are depicted in representative volumes of interest (Fig. S1). CNO or vehicle were administered at time 0; Hp: hippocampus; mPFC: prefrontal cortex; DRN: dorsal raphe nucleus; NAc: nucleus accumbens; Th: thalamus; Hyp: hypothalamus; SS: somatosensory cortex; Cb: cerebellum.

**Figure S3.**
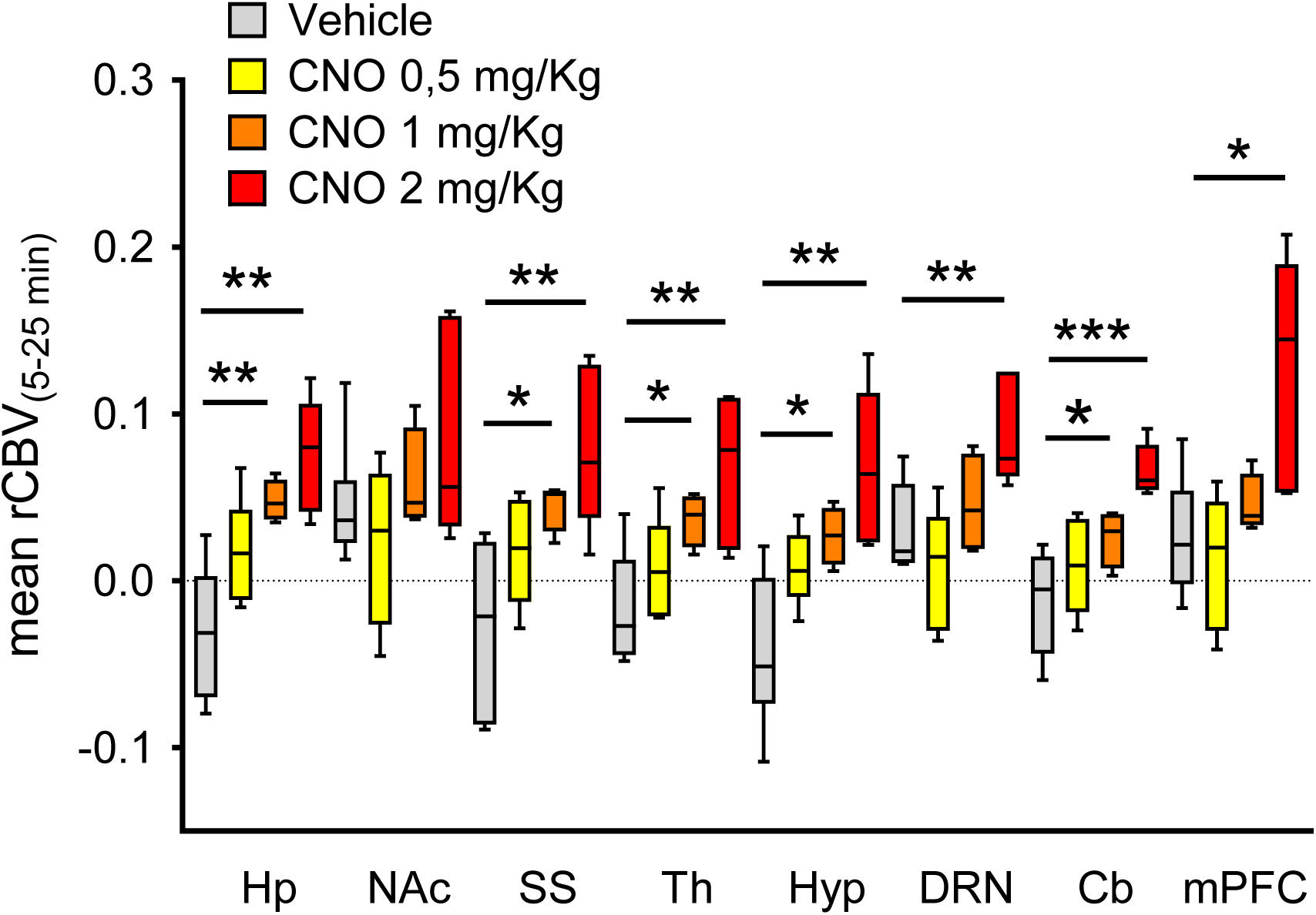
Quantification of the mean rCBV response elicited by the three doses of CNO tested in this study (0.5, 1 and 2 mg/kg i.v.) in representative volumes of interest (*p<0.05, **p< 0.01, Student t test, false discovery rate correction q=0.05). Hp: hippocampus; NAc: nucleus accumbens; SS: somatosensory cortex; Th: thalamus; Hyp: hypothalamus; DRN: dorsal raphe nucleus; Cb: cerebellum, mPFC: prefrontal cortex.

**Figure S4.**
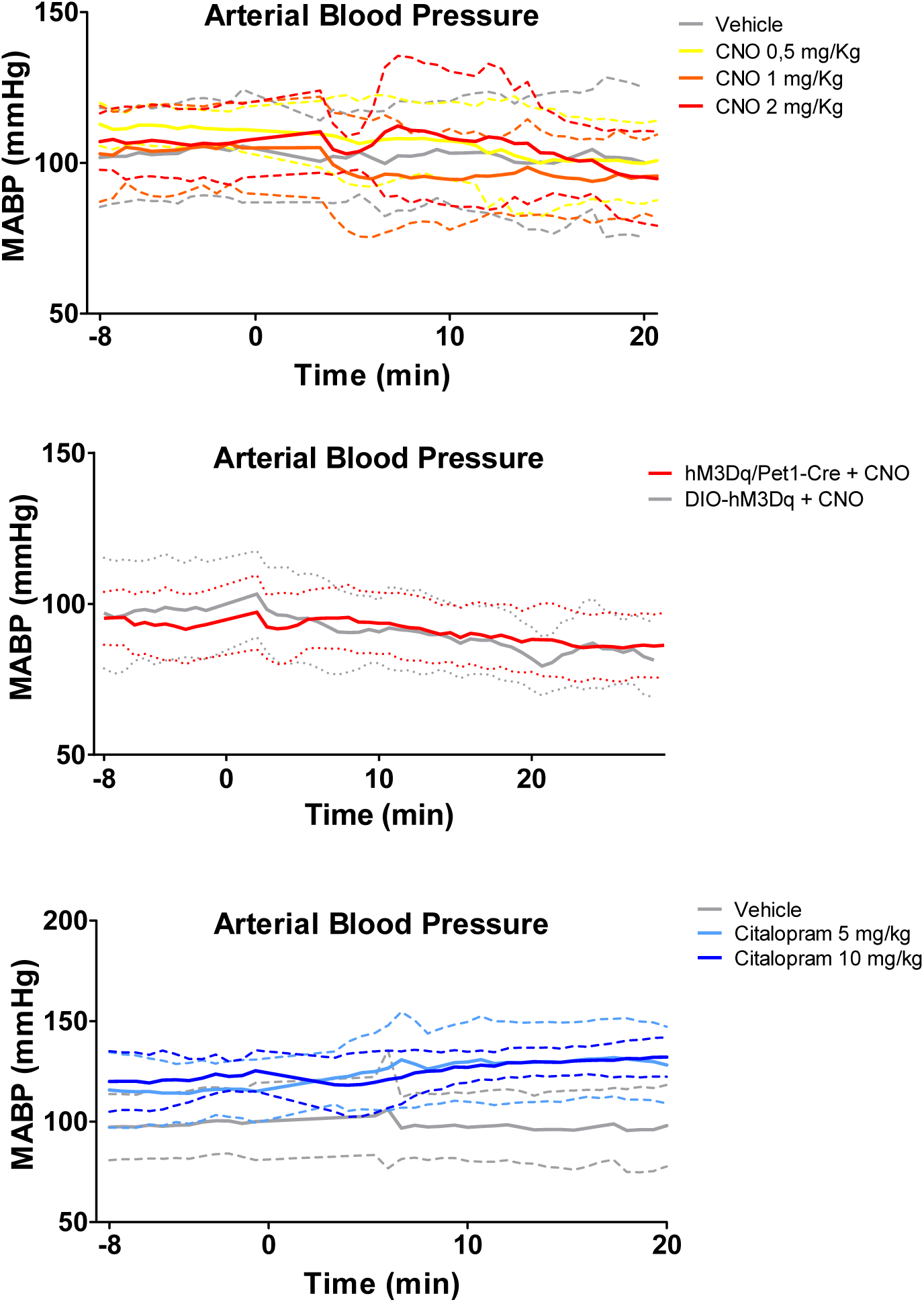
Temporal profile of mean arterial blood pressure (MABP). (A) MABP timecourse upon vehicle (n = 5) or CNO administration at doses of 0.5 (n=6), 1 (n=4) or 2 mg/kg i.v. (n=5). (B) MABP upon CNO administration (0.5 mg/kg i.v.) to hM3Dq/Pet1-Cre (n=20) or DIO-hM3Dq (n=14) control subjects. MABP timecourse (C) upon vehicle (n=10) or citalopram administration at dose of 5 (n=8) or 10 mg/kg i.v. (n=7) to wild type mice. Data are plotted as mean±SD within each group, drugs and vehicles were administered at time 0.

**Figure S5.**
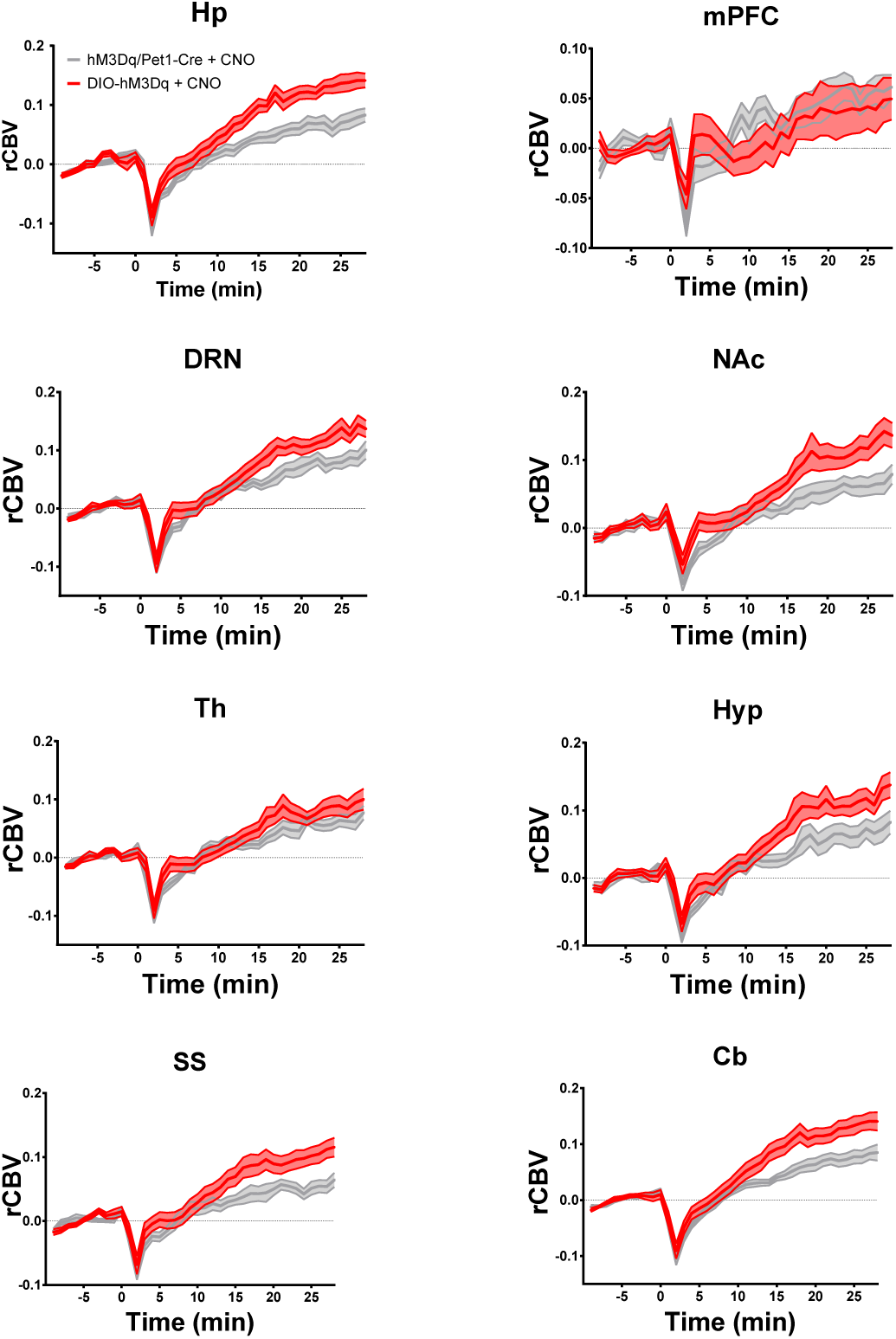
rCBV response produced by intravenous CNO administration to hM3Dq/Pet1-Cre (n=20) and DIO-hM3Dq controls (n=14) in representative volumes of interest. CNO was administered at a dose of 0.5 mg/kg i.v. to both experimental groups. CNO was administered at time 0; Hp: hippocampus; mPFC: prefrontal cortex; DRN: dorsal raphe nucleus; NAc: nucleus accumbens; Th: thalamus; Hyp: hypothalamus; SS: somatosensory cortex; Cb: cerebellum.

**Figure S6.**
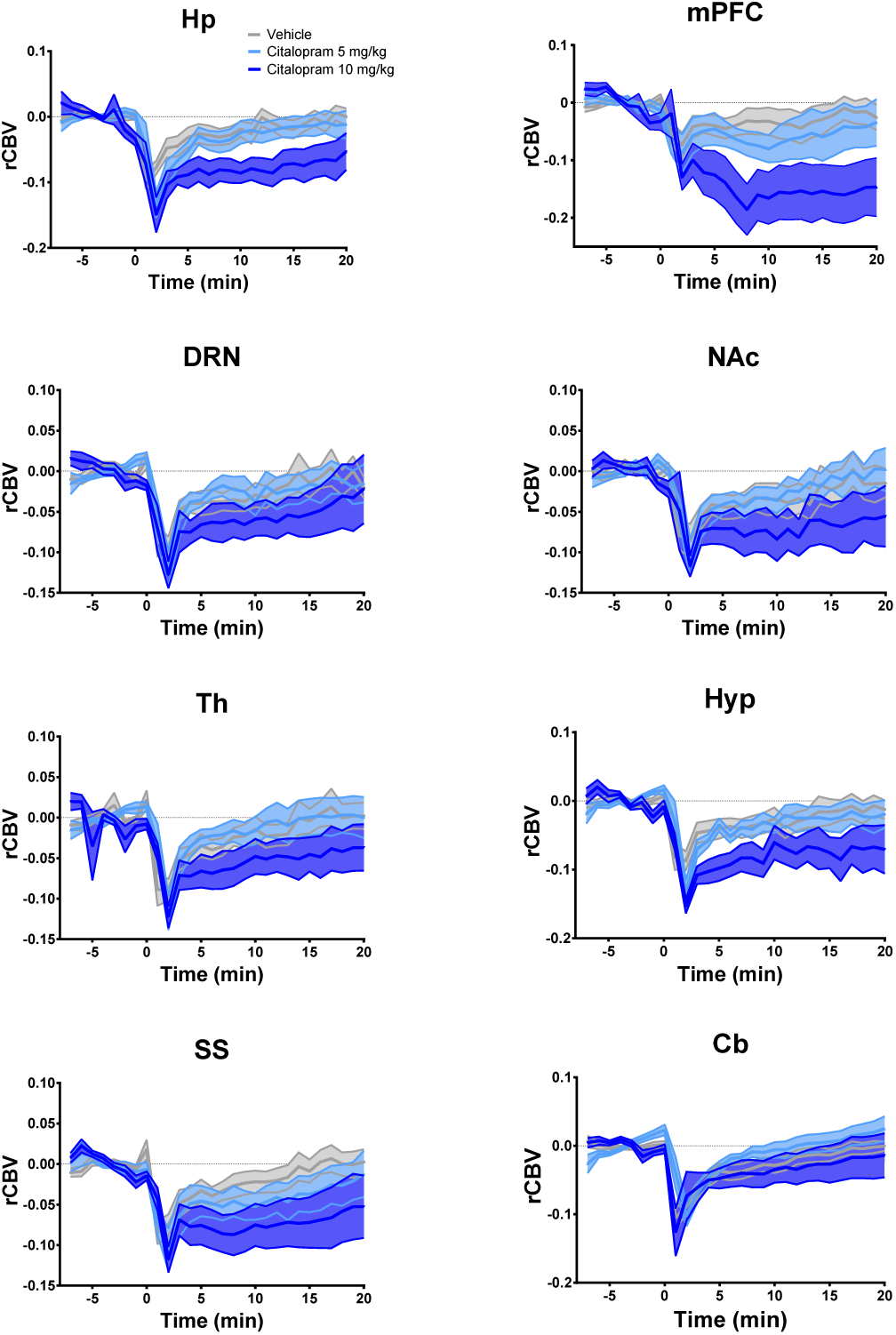
rCBV response produced by intravenous citalopram or vehicle administration in wild-type mice. Citalopram was administered at a dose of 5 (n=8) or 10 mg/kg i.v. (n=7). Citalopram or vehicle (n=10) were administered at time 0; Hp: hippocampus; mPFC: prefrontal cortex; DRN: dorsal raphe nucleus; NAc: nucleus accumbens; Th: thalamus; Hyp: hypothalamus; SS: somatosensory cortex; Cb: cerebellum.

## MATERIAL & METHODS

### Ethical statement

All research involving animals were carried out in accordance with the European directive 86/609/EEC governing animal welfare and protection, acknowledged by the Italian Legislative Decree no. 116, 27 January 1992. Animal research protocols were also reviewed and consented to by a local animal care committee.

### Mouse line generation

To enable stable and reproducible pan-neuronal stimulation of 5-HT producing cells, we first generated a conditional knock-in hM3Dq mouse line. The hM3Dq sequence in-frame-fused with the mCherry reporter and flanked by two couples of Lox sites (LoxP and Lox2722, DIO-hM3Dq, Double-floxed Inverse ORF) was isolated by EcoRI digestion from pAAV-hSyn-DIO-hM3Dq construct (a kind gift From Dr. Bryan Roth) and inversely cloned downstream to the CAG promoter in the pCX-CAG-eGFP plasmid, thus replacing the eGFP coding sequence (CDS)(Krashes *et al*, 2011). The CAG-DIO-hM3Dq fragment was extracted from the pCX-CAG-hM3Dq by a SpeI/XbaI double digestion and cloned into the unique XbaI site placed (Krashes *et al*, 2011)between the right (RA) and the left (LA) ROSA26 homology arms included in the pROSA26-1^Sor^ plasmid. Pgk-Neo/Kana (NEO) and Pgk-Difteric Toxin A cassettes (DTa) were used for positive and negative selection, respectively. PvuI-linearized pROSA26-1^Sor^-LA-CAG-DIO-hM3Dq-NEO-RA-DTa was electroporated in E14Tg2a.4 embryonic stem cells (ESCs) as previously described (Pelosi *et al*, 2015). Positive recombinants identified by southern blot were microinjected in the host C57BL/6J blastocysts, which were injected in utero of pseudopregnant CD1 females (3.5 dpc) to give rise to chimeras (n=29). Animals were routinely genotyped by PCR DNA amplification with specific oligonucleotides as primers: 5’-GAGGGGAGTGTTGCAATACC-3’ as forward, and 5’-AGTCTAACTCGCGACACTGTA-3’ as reverse, for the wild-type ROSA26 allele; the alternative reverse 5’-GTCCCTATTGGCGTTACTATG-3’ for the DIO-hM3Dq allele; 5’-GTCATCTCCTTTGTCCTTTGG-3’ as forward and 5’-GGAGCTGGGTTTCCAGCTC-3’ as reverse for the hM3Dq CDS. Before chemo-fMRI experiments, animals were backcrossed on C57BL/6J for a least six generations. To enable the expression of hM3Dq in 5-HT producing neurons, mice were then crossed with Pet1_210_-Cre mice (Barrett *et al*, 2016; Pelosi *et al*, 2014). The resulting hM3Dq^+/-^/Pet1-Cre mice were image during young adulthood (12-20 weeks). Control studies with CNO and citalopram were carried out using male adult (12-20 weeks) C57BL/6J mice.

### Immunohistochemical analyses

Mice were deeply anesthetized with avertin 1.25 % and perfused transcardially with 4% paraformaldehyde. Brains were dissected, post-fixed overnight in 4% paraformaldehyde at 4°C and sectioned in slices 50 μM thick by a Leica Microsystems vibratome. In mCherry and GFP immunodetection studies, free floating sections were incubated at 4°C overnight with rabbit anti-RFP (ab62341, Abcam, 1:500) and chicken anti-GFP (Ab13970, Abcam, 1:1000). Sections were then incubated overnight at 4°C with goat anti-rabbit (Rhodamine Red, Invitrogen, 1:500) and goat anti-chicken (Alexa Fluor 488, Invitrogen, 1:500). Animals employed for c-Fos mapping (study #4, described below) were euthanized ninety minutes after CNO administration and processed as described above for immunohistochemical mapping. Free-floating sections were incubated overnight at room temperature with goat anti-cFos (SC-52-G, Santa Cruz Biotechnology, 1:1000) in a 5% horse inactivated serum solution. Sections were then incubated overnight at 4°C with donkey anti-goat (Alexa Fluor 488, Invitrogen, 1:500). Cell counts for c-Fos quantification and mCherry/eGFP co-localization analysis were performed using FIJI-ImageJ.

### Electrophysiology

Brains of hM3Dq^+/-^/Pet1-Cre were sectioned to obtain 300 μm coronal slices encompassing the dorsal raphe nucleus. Slices were submerged in normal, oxygenated aCSF (28-30° C, 2mL/min flow rate) for at least 30 minutes before performing whole-cell patch clamp experiments. Borosilicate electrodes with a pipette filled with internal solution (135 KMeSO4, 10 KCl, 10 HEPES, 1 MgCl2, 2 Na2-ATP, 0.4 Na3-GTP (pH 7.2-7.3, 280-290 mOsm/kg). were used to patch cells in the DRN. Signals were acquired using a Multiclamp 700B amplifier and analyzed with Clampfit 10.3 software (Molecular Devices, Sunnyvale, CA, USA). The effects of CNO were determined in current clamp mode. After 5 minutes of stable baseline, CNO (10 μM) was bath applied for 10 minutes while recording changes in membrane potential. For excitability experiments, the current threshold (rheobase) necessary to induce cell firing were determined in current clamp mode using a current ramp protocol from 0 to 100 pA. Next, a 10 pA current step protocol from 0 to 200 pA was applied, from which V-I plots were determined (i.e., the number of action potentials vs. current).

## Drug formulation and pharmacological treatments

All drugs were administered intravenously. We chose this route of administration to maximize the rate of fMRI signal change, with the aim to induce sharp fMRI responses that can be more easily discriminated from linear alterations in baseline due to scanner, contrast agent elimination or physiological drifts. Clozapine-n-Oxide (Sigma Aldrich) was dissolved in saline solution at a concentration of 0,125 μg/ml. The CNO dose employed for chemo-fMRI mapping (0,5 mg/kg i.v., volume 10 mL/kg) was selected out of dose-response experiments (0,5-1-2 mg/kg i.v.) described in the result section. Citalopram (Sigma Aldrich) was dissolved in saline solution at a concentration of 3 or 1.5 ml/kg for 10 mg/kg and 5 mg/kg dosing, respectively. The citalopram doses tested were previously shown to be behaviorally effective in C57Bl/6J mice (Browne and Fletcher, 2016).

Pilot c-Fos mapping studies in which we employed fMRI dose regimen of CNO, showed that the prolonged restriction required for intravenous administration of CNO results in broad unspecific FOS increases with respect to baseline conditions. We therefore opted for the administration of a dose of 2 mg/kg intraperitoneally to account for the reduced maximum plasma concentration associated with intraperitoneal administration with respect to intravenous administration. The dose employed is in line with the amounts of CNO tested by other investigators in DREADD-based studies (Roth, 2016).

### Functional Magnetic Resonance Imaging (fMRI)

Animal preparation for functional magnetic resonance imaging has been previously described in great detail (Dodero *et al*, 2013). The protocol utilized is optimized for physiological stability and permits monitoring of peripheral parameters critical to the success of pharmacological fMRI (phMRI), such as peripheral blood pressure (Ferrari *et al*, 2012) and arterial blood gases. Briefly, mice were anaesthetized with isoflurane (5% induction), intubated and artificially ventilated (2.5% surgery). The left femoral artery was cannulated for continuous blood pressure monitoring and blood sampling. Surgical sites were infiltrated with tetracaine, a non-brain-penetrant local anesthetic (Ferrari *et al*, 2010). At the end of surgery, the animal was placed in supine position onto a water-heated custom cradle and isoflurane was discontinued and replaced by halothane (0.7%), an anesthetic that preserves cerebral blood flow auto-regulation and neurovascular coupling (Gozzi *et al*, 2007). Functional data acquisition started 30 minutes after isoflurane cessation. Ventilation parameters were adjusted to maintain arterial p_a_CO_2_ levels < 40 mmHg (Pepelko and Dixon, 1975) and p_a_O_2_ levels > 90 mmHg, values which correspond to the 98% of hemoglobin saturation (Supplementary table I).

fMRI data were acquired as previously described (Squillace *et al*, 2014) on a 7T Pharmascan (Bruker, Ettlingen, Germany) by using a 72-mm birdcage resonator and a 4 channel anatomical shaped Bruker mouse brain coil, placed dorsally to the animal head. Co-centered anatomical and fMRI images were acquired using a Rapid Acquisition Relaxation-Enhanced and a Fast Low-Angle Shot MRI sequence (TReff = 288 ms, TEeff = 3.1 ms, α=30°; 180 x 180 x 600 μm resolution, dt = 60 s, Nr = 60 corresponding to 60 min total acquisition time). Images were sensitized to reflect alterations in relative cerebral blood volume (rCBV) by previous administration of 5 μl/g of blood-pool contrast agent (Molday Ion, Biopal, Worcester, MA, USA). Twenty-five minutes later each subject received an intravenous administration of vehicle or drug. fMRI responses were mapped and quantified as previously described (Galbusera *et al*, 2017). Briefly, fMRI time series were spatially normalized to a common reference space and signal intensity changes were converted into fractional rCBV changes. rCBV time series before and after drug or vehicle injections were extracted and analyzed. Voxel-wise group statistics was performed using FEAT Version 5.63, with 0.5 mm spatial smoothing and using a boxcar input function that captured the main alterations in rCBV signal observed upon pharmacological challenge with respect to vehicle baseline in representative volumes of interest (Figure S1). Specifically, the chemo-fMRI response to 5-HT stimulation was used using a boxcar function covering a 10-28 min post-injection time window, while citalopram was mapped using a boxcar regressor depicting a 0-25 min post injection time window.

The composition of experimental groups was as follows:

Study 1: – CNO dose-response in C57Bl6J mice: intravenous treatment with CNO (0,5 mg/kg n=6; 1 mg/kg n=4; 2 mg/kg n=5) or vehicle (n=6).

Study 2: Chemo-fMRI of 5-HT neurons: intravenous treatment with CNO in hM3Dq/Pet1-Cre (*n* = 20), or control DIO/hM3Dq (n = 14).

Study 3: fMRI of citalopram in C57Bl6J mice: intravenous injection of citalopram (5 mg/kg n=8; 10 mg/kg n=7) or vehicle (n=10).

Study 4: CNO-induced FOS-mapping in hM3Dq/Pet1-Cre (n = 5) or control DIO/hM3Dq mice (n = 5).

